# Spontaneous spiking statistics form unique area-specific fingerprints and reflect the hierarchy of cerebral cortex

**DOI:** 10.1101/2025.05.13.653871

**Authors:** Aitor Morales-Gregorio, Robin Gutzen, Sofia Paneri, Panagiotis Sapountzis, Alexander Kleinjohann, Sonja Grün, Alexa Riehle, Xing Chen, Thomas Brochier, Georgia G. Gregoriou, Bjørg E. Kilavik, Sacha J. van Albada

## Abstract

The cerebral cortex is hierarchically organised from sensory to higher cognitive areas^1–4^. Several dynamical^5–8^ and anatomical^1–4,8^ measures, such as timescales and neurotransmitter receptor expression, have independently been linked to the cortical hierarchy. However, a systematic and quantitative characterisation of the relationship between spontaneous spiking activity and the cortical hierarchy remains elusive. Here, we test the hypothesis that single-neuron spontaneous spiking statistics uniquely characterise each cortical area, and that they quantitatively correlate with the cortical hierarchy. We study the spontaneous activity of neurons in seven macaque cortical areas (V1, V4, DP, 7A, M1, PMd, PFC)^9–12^ in the eyes-open and eyes-closed conditions recorded in a dim-lit room. First, we uncover that the firing rate, inter-spike interval variation, and cross-correlation form a unique fingerprint of the cortical areas, but only when considering them in combination. Second, we show that the differences between the spiking statistics correlate with multiple anatomical markers^1,2,4,13–17^ of the cortical hierarchy. This effect is much stronger in the eyes-closed condition, suggesting that visual input or the expectation thereof modulates the hierarchical organisation of spontaneous activity. We also observe an increase in timescales up the hierarchy, in agreement with previous findings^5,18,19^. In conclusion, we demonstrate that spontaneous single-neuron spiking activity reflects the hierarchical organisation of the cerebral cortex: distinct spiking statistics for hierarchically distant areas; similar statistics for nearby areas. Our results thus add a new dynamical dimension to the concept of the cortical hierarchy.

## Introduction

Information in the cerebral cortex is processed in multiple stages, following a hierarchy^1^. Within this hierarchy, lower-order regions are thought to handle localized sensory inputs while higher-order regions integrate signals into abstract representations. Several anatomical features, such as structural connectivity^1–3,20^, neuron^13,14^ and receptor^4,15,21^ densities, spine counts^16,22^, and myelination^17^, reflect the cortical hierarchy. All these anatomical markers of the hierarchy are known to strongly correlate with each other^4,15^, suggesting a complex relationship between the structural anatomy and information processing along the hierarchy.

Neural activity also varies along the cortical hierarchy. For example, inter-spike interval (ISI) variation^23^ of awake behaving animals differs significantly across areas at different hierarchical levels and is surprisingly consistent across mammals for the same area^6^. Functional connectivity^24,25^ and visual response measures^7^ have been reported to align with the hierarchical organisation. Autocorrelation timescales also follow a hierarchical gradient in macaques^5,26–28^, humans^18,27,29,30^, rats^28^, and less clearly in mice^7,19,27,28^. While prior studies have independently examined anatomical markers and neural dynamics, the systematic relationship between these features remains unexplored. Specifically, no study reports systematic quantitative correlation between spontaneous spiking activity at the single-neuron level and cortical hierarchy across many areas in primates.

Here, we report evidence supporting the hypothesis that cortical spontaneous activity forms a unique fingerprint of each area and follows an anatomical hierarchical gradient. We use extracellular electrophysiological data from seven cortical areas (V1, V4, DP, 7A, M1, PMd, PFC) in several macaques^9–12^ (*N* = 6, *Macaca mulatta*), and measure single-neuron statistics; i.e. firing rate, ISI variation, and the mean and standard deviation of cross-correlations of individual neurons with all other simultaneously recorded neurons. We study spontaneous activity because it is expected to be less biased toward external stimuli and more closely related to the underlying anatomical structure than evoked activity. We test whether the spontaneous activity varies across cortical areas, and find significant differences in the spontaneous spiking statistics across all areas, especially when the statistics are considered multivariately. These differences strongly correlate with several anatomical markers of the cortical hierarchy: neuron densities^13^, receptor densities^4^, spine counts^16^, and hierarchical levels estimated from laminar cortico-cortical connectivity^2^. Moreover, the correlation between dynamics and anatomy is stronger during eyes-closed periods. We further confirm^5,26–28^ that autocorrelation timescales also exhibit a consistent correlation with the hierarchy. All in all, the results suggest a key role for cortical anatomy in shaping spontaneous spiking activity and add a new dynamical dimension to the concept of cortical hierarchies.

## Results

### Multivariate spiking statistics of spontaneous activity

Does spontaneous neural activity systematically vary across cortical areas? To answer this question, we studied spontaneous intracortical electrophysiology recordings from various experiments in macaques^9–12^. The data were recorded from seven areas along the visuomotor hierarchy (V1, V4, DP, 7A, PFC, PMd, and M1). Approximate recording locations are shown in Figure 1a. The data were spike sorted based on their waveforms to retain only single-unit activity (SUA), after which any SUA suspected of originating from hardware noise or cross-talk was excluded^31^ (Figure 1b). The different experiments varied in recording length (Figure S4a,b), hence we split the data into 5-s segments to allow for unbiased comparisons. To reduce nonstationarities, segments were only included if the monkey remained in the same behavioural state (eyes-open or eyes-closed) throughout the whole segment. See Electrophysiological data collection of spontaneous activity for details on the data collection and preprocessing.

**Figure 1:**
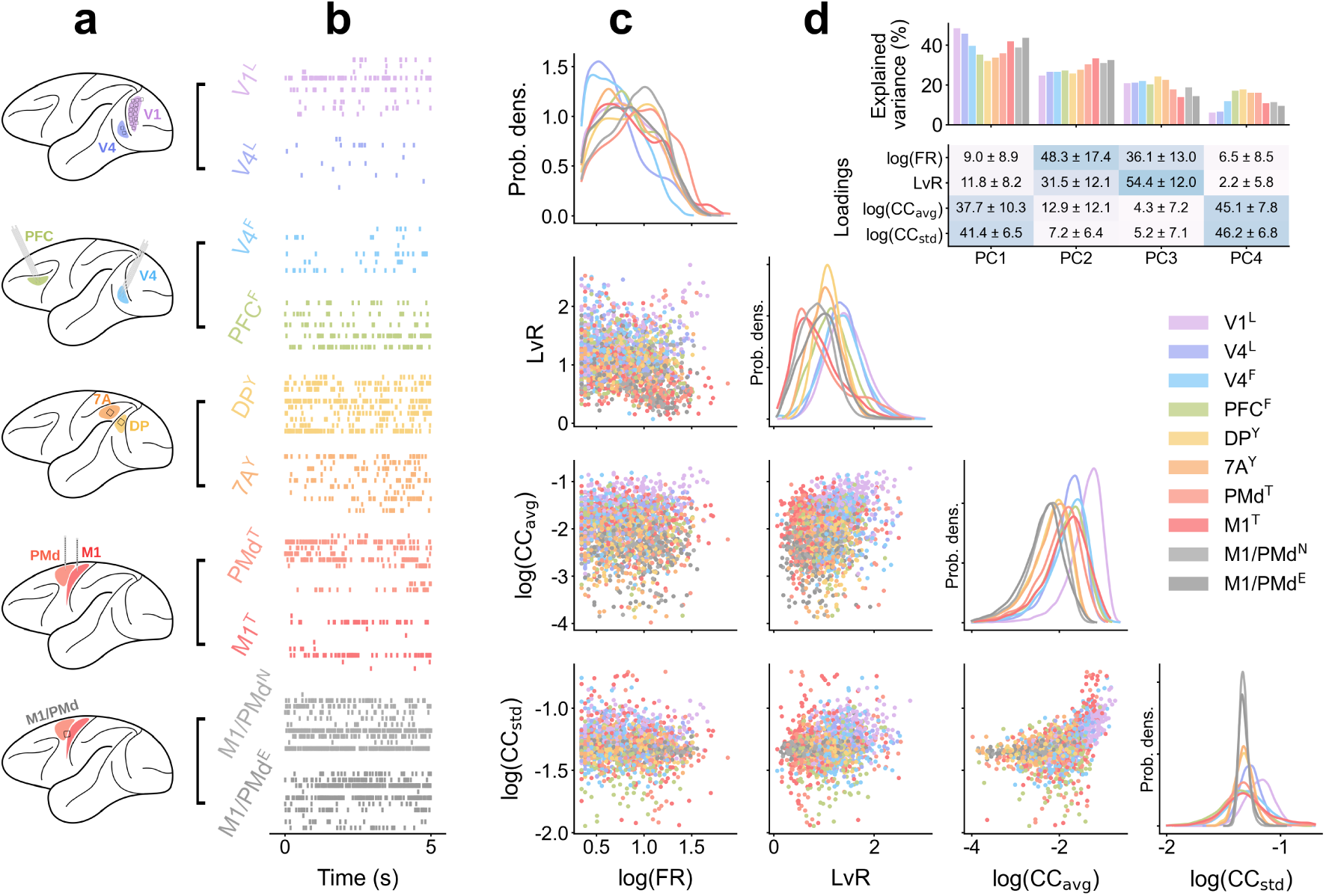
Sample of available experimental data and their spontaneous spiking statistics. Same color scheme across panels. **a)** Schematic representation of the data recording locations. **b)** Sample recordings of simultaneous spike trains from each dataset for a 5-s segment; superscripts indicate the subject identity. **c)** Pair plot of single-neuron spiking statistics. Panels along the diagonal show the kernel density estimates (KDE) of individual distributions, whereas off-diagonal panels show scatter plots where each point is calculated from a 5-s spike train from one neuron. **d)** (Top) Variance explained by each of the four principal components (PC) of the multivariate spiking statistics. (Bottom) Loadings of each spiking statistic (average across datasets ± standard deviation). PCA performed separately for each dataset.

To characterize the spiking activity, we calculated single-neuron statistics for each data segment and assembled them into a multivariate point cloud (Figure 1c). The statistics were firing rate (FR), revised local variation (LvR) of inter-spike intervals^23^, and the mean and standard deviation of each neuron’s cross-correlation with all other neurons in the same area and subject (CC_avg_ and CC_std_). The LvR measures the irregularity of spike trains by considering triplets of subsequent spikes, thus reducing the dependence on firing rate fluctuations. The cross-correlation statistics are related to the dimensionality of the population activity^32–36^. Since the FR, CC_avg_ and CC_std_ had highly skewed distributions, we took their base-10 logarithm, rendering the distributions more symmetric. See Statistics of spiking neuronal data for details.

Note that we did not include the single-neuron timescales here, because they are difficult to estimate from short recordings^37–39^, as we illustrate in Figure S4. Instead, we separately studied timescales based on longer segments (see Autocorrelation timescales also reflect hierarchy).

We visualized the four-dimensional cloud of single-neuron statistics (Figure 1c) as a pair plot, where each subplot displays two statistics, and the diagonal subplots show the kernel density estimates (KDE) of individual distributions. The spiking statistics for each animal and area are shown individually in Figure S1–S3. To assess whether our chosen statistics were relevant and linearly independent, we estimated the explained variance and loadings using PCA. The principal components each explained a substantial proportion of variance (at least 5–20%, (Figure 1d top), demonstrating that all the selected spiking statistics capture independent aspects of neural dynamics.

The experiments were highly heterogeneous, with different subjects, hardware setup, neuron count, and recording duration. Calculating the multivariate distribution of single-neuron spontaneous spiking statistics allows comparison of heterogeneous datasets by summarizing them in a common space.

### Multivariate spiking statistics form a unique fingerprint of each cortical area

Visual inspection reveals apparent differences between the distribution of spiking statistics across brain areas (Figure 1c). We test whether these differences are statistically significant. The inclusion of multiple spiking statistics provides a comprehensive characterization of each area, since each statistic captures some unique information about each area. Hence, we expected the joint distributions of multiple statistics to show more differences between areas than each single statistic on its own. For example, while FR may not always uniquely characterise each area, the combination of all four statistics could. We tested this further hypothesis by testing whether the means of the distributions of spiking statistics differed significantly across areas and subjects. First, we performed a pairwise univariate analysis of variance (ANOVA) for each of our four statistics (Figure 2a–d). Second, we performed pairwise multivariate ANOVA (MANOVA) tests (Figure 2e), where all four spiking statistics were considered in combination. See Statistical testing with ANOVA and MANOVA for details about the tests. The univariate ANOVA tests (Figure 2a–d) revealed that no single spiking statistic consistently differentiates activity across all areas. However, when all the spiking statistics were aggregated and tested as an ensemble with the MANOVA test (Figure 2e) nearly all dataset pairs became statistically different. The only exception was the MANOVA test between V4^F^ ↔ V4^L^, for data originating from the same area in two different subjects, recorded with different hardware, and in different laboratories. This suggests that the observed differences are not explained by differences between subjects, hardware, or laboratories.

**Figure 2:**
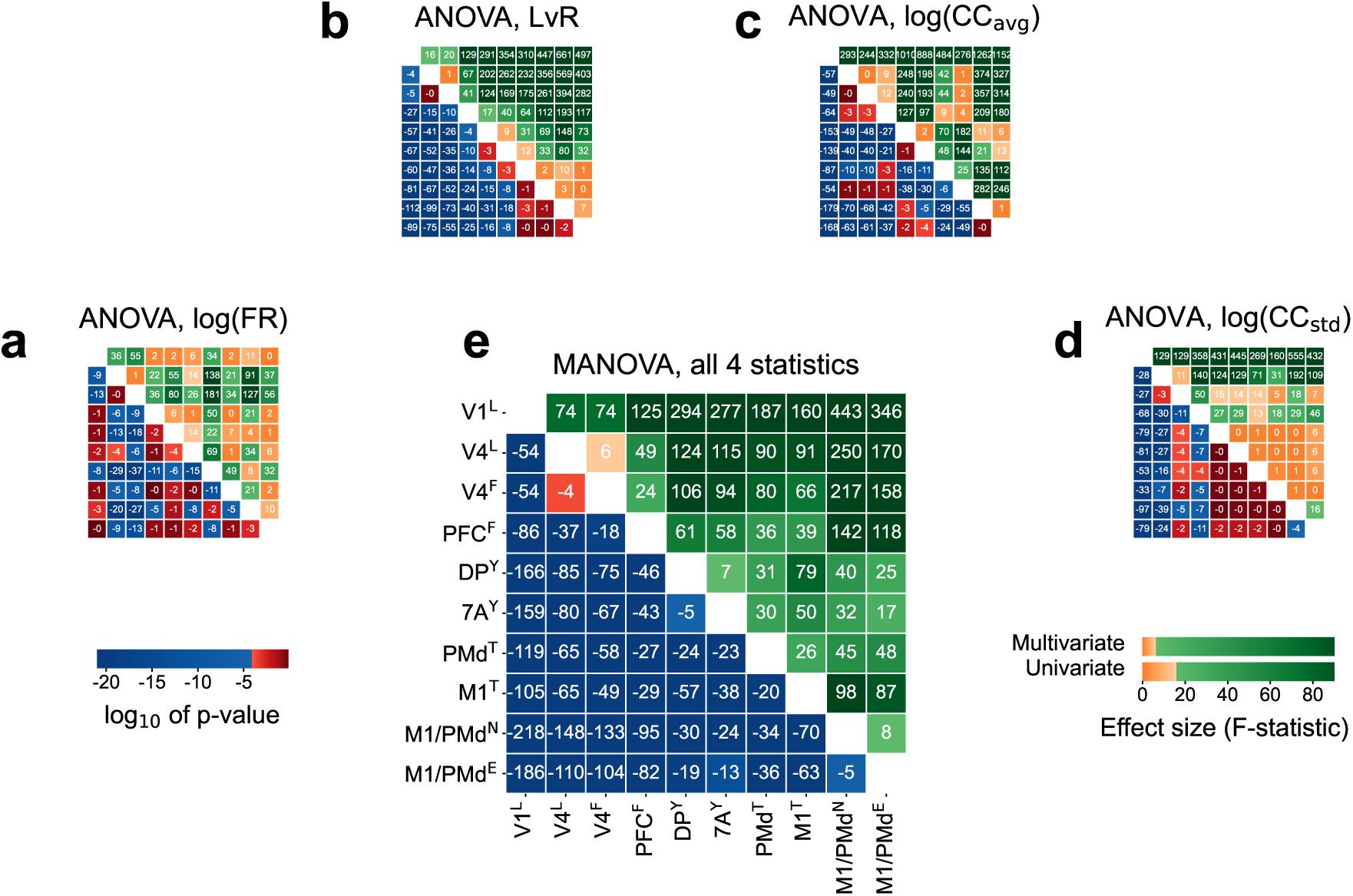
Uni- and multivariate pairwise comparisons of spontaneous spiking statistics. We test the null hypothesis that two or more datasets have the same mean. In all panels, lower triangular entries show the base-10 logarithm of the p-values and the upper triangular entries show the F-statistic. Significance levels (*α* = 0.05*/k*) are corrected for multiple testing following the Bonferroni correction (*k* = 630, including the tests from Figure S8a). In all panels, the significance level is color-coded, with red p-values (orange F-statistics) denoting non-significant results. **a-d)** Univariate pairwise analysis of variance (ANOVA) tests. **e)** Multivariate pairwise analysis of variance (MANOVA) tests.

The statistical tests support our hypothesis that the spiking statistics are area-specific, but only if considered multivariately. Thus, the multivariate spiking statistics form a dynamical fingerprint of each cortical area.

### Dynamical dissimilarity reveals hierarchical organisation of cortical areas

The statistical tests so far revealed that the multivariate spiking statistics differ between areas, but not how similar or different they are. To quantify those differences, we introduce the dynamical dissimilarity, which we define as the Wasserstein distance (WS) between the multivariate spiking statistics of two areas (Figure 3a). We chose to use the Wasserstein distance because it is symmetric, non-parametric, and numerically robust on point clouds. The dynamical dissimilarity takes a positive value between zero (for identical distributions) and infinity, and can be applied to compare any two point distributions both univariately or multivariately. Low values of the dynamical dissimilarity indicate high similarity. We calculated all within- and across-area dynamical dissimilarities (Figure 3a), and found that the within-area dissimilarities were consistently smaller than across-area dissimilarities (Figure 3a inset). See Dynamical dissimilarity for comparison of multivariate and univariate distributions for a detailed description.

**Figure 3:**
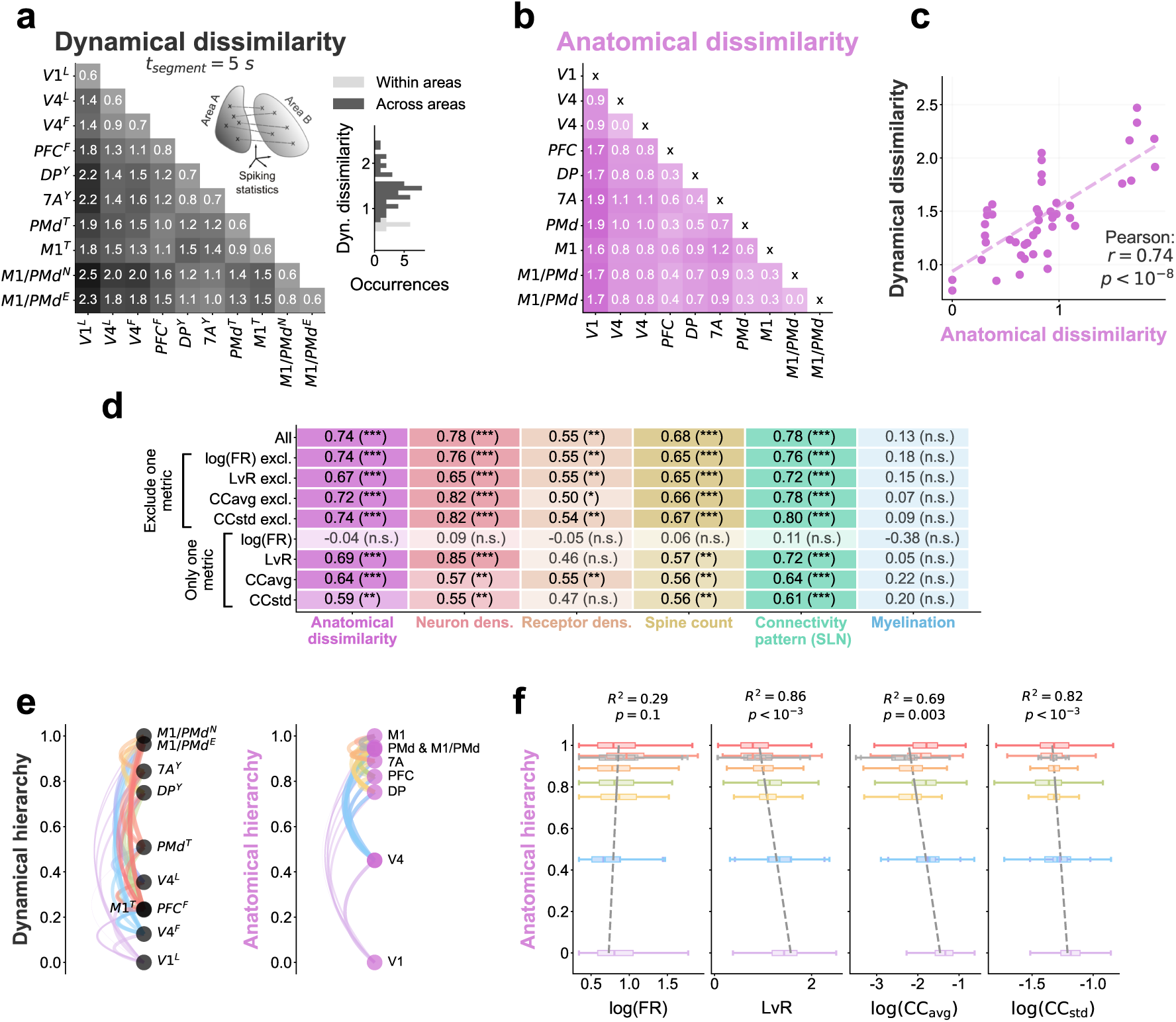
Dynamical dissimilarity between spiking activity statistics strongly correlates with anatomical markers of cortical hierarchy. **a)** Dynamical dissimilarity between all datasets. The inset diagram schematically illustrates the Wasserstein distance (WS). The inset histogram shows the distribution of dynamical dissimilarity within and across datasets. **b)** Anatomical dissimilarity between all areas, defined from the combination of neuron density, receptor density, spine counts, connectivity patterns, and myelination. **c)** Dynamical dissimilarity strongly correlates with anatomical dissimilarity; Pearson correlation coefficient and corresponding t-test p-value are shown. **d)** Pearson r between anatomical dissimilarity and modified versions of the dynamical dissimilarity: excluding one metric at a time (leave-one-out) or only including a single metric. Stars indicate result of t-test: *p >* 0.05*/k* (n.s.), 0.05*/k > p >* 0.01*/k* (*), 0.01*/k > p >* 0.001*/k* (**), *p <* 0.001*/k* (***). Bonferroni corrected for multiple testing (*k* = 61, also including the tests from Figure 5d). **e)** Estimates of the hierarchy from the dynamical and anatomical dissimilarities. **f)** Boxplot of the spiking statistics for each area compared to their position in the anatomical hierarchy. Weighted least squares (WLS) fit, its coefficient of determination *R*^2^ and t-test p-value are shown.

Moreover, the dynamical dissimilarities were highly structured (Figure 3a). They were smallest for neighbouring (7A ↔ DP or PMd ↔ M1) or equivalent areas in different subjects (M1*/*PMd^N^ ↔ M1*/*PMd^E^ or V4^F^ ↔ V4^L^), and largest between functionally distant areas (e.g. V1 ↔ M1*/*PMd). This trend suggests a hierarchical organisation of dynamical differences.

To test the hierarchical organisation of dynamical dissimilarities we compared them to inter-area differences in several known anatomical markers of the hierarchy: neuron densities^13^, receptor densities^4^, spine counts^16^, hierarchical levels estimated from laminar cortico-cortical connectivity^2^, and myelination^17^. See Anatomical data for hierarchical characterization for a detailed description of the anatomical data.

To relate the dynamical dissimilarities to the anatomical dissimilarities, we calculated the Euclidean distance between the 5D data points representing the anatomical markers for each pair of areas (Figure 3b). Each marker was normalized between 0 and 1 prior to measuring the anatomical dissimilarity. We found a strong correlation (Pearson’s *r* = 0.74) between the dynamical and anatomical dissimilarities (Figure 3c).

The receptor densities were available for 14 different receptor types, but we included only their average in our anatomical dissimilarity, so as not to eclipse all other anatomical markers. We also tested the individual receptor densities and most of them strongly correlated with the full dynamical dissimilarity (Figure S5).

To analyze the individual contributions of each spiking statistic, we compared the anatomical dissimilarity to the dynamical dissimilarity when one metric was excluded (’Exclude one metric’ in Figure 3d) and also to the dynamical dissimilarity when a single spiking statistic was included (’Only one metric’ in Figure 3d). Furthermore, we extended the comparisons to each individual anatomical marker (Figure 3d); corresponding scatter plots are shown in Figure S6. This analysis reveals which spiking statistic relates most to which anatomical feature.

Notably, log(FR) did not correlate with any anatomical markers when considered univariately. Furthermore, myelination^17^ did not correlate with any of the spiking statistics. If myelination is excluded from the anatomical dissimilarity the correlation in Figure 3c increases slightly (*r* = 0.78*, p <* 10*^−^*^9^).

We embed the dynamical and anatomical dissimilarities into a 1-dimensional line, representing the hierarchy (see Estimation of hierarchies from dynamical and anatomical dissimilarities, Figure 3e). A similar (but not equal) area ordering emerged from both dynamics and anatomy (Figure 3e).

To study how dynamics change along the hierarchy we compared the mean of each spiking statistic against the anatomical hierarchy using Weighted Least Squares (WLS), weighted by the inverse of the variance of each statistic (Figure 3f). No significant trend was found for log(FR). The WLS indicated that LvR decreases along the hierarchy: the spike trains in V1 were irregular (high LvR), and the spike trains in motor areas more regular (lower LvR). Furthermore, the WLS showed that log(CC_avg_) and log(CC_std_) decrease along the hierarchy, suggesting that the dimensionality^32–36^ of the population activity decreases along the hierarchy.

In conclusion, the spontaneous spiking statistics robustly correlate with anatomical markers of the cortical hierarchy, strongly supporting our hypothesis that cortical spontaneous activity follows an anatomical hierarchical gradient.

### Correspondence between dynamics and anatomy is strongest with the eyes closed, and nearly disappears with the eyes open

The results until this point combined all behavioural states. However, spiking activity is known to vary between different behaviours^36,40^. We expect stronger external inputs during the eyes-open (EO) than eyes-closed (EC) periods, which may affect the global ongoing activity of the cortex. During EC, the activity may more closely reflect the local anatomical structure of each area, due to the reduced external stimuli. Analyzing the differences between EO and EC thus provides potential information about the relative contributions of local and global factors to the observed hierarchical organisation. We therefore split the activity by behaviour and study the changes in the dynamical dissimilarity (Figure 4).

**Figure 4:**
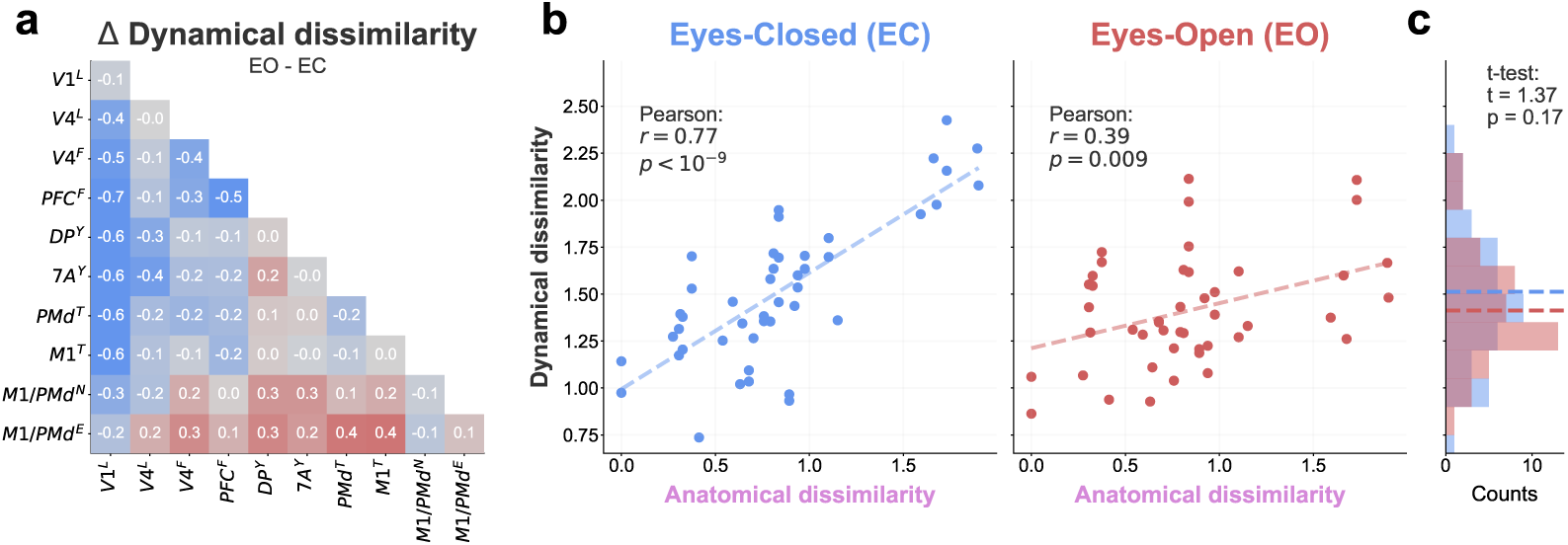
Dynamical dissimilarity changes between eyes-open (EO) and eyes-closed (EC) epochs. **a** Difference between the EO and EC dynamical dissimilarity. Negative values (blue) indicate higher dissimilarity in EC, conversely positive values (red) indicate higher dissimilarity in EO. **b** Correlation between dynamical and anatomical dissimilarities in EC and EO. The correlation is much weaker for EO. **c** Histograms of the dynamical dissimilarity distributions for EO and EC. Dashed lines indicate the mean value. The EO and EC distributions are not statistically different (two-sided t-test).

First, we measured the dynamical dissimilarities for samples during eyes-open (EO) or eyes-closed (EC) periods. We found that visual areas, especially V1, have higher dynamical dissimilarity in EC than EO; whereas motor areas have higher dissimilarities in EO than EC (Figure 4a). Second, we compared the correlation between dynamical and anatomical dissimiliaries for the EC and EO conditions (Figure 4b), revealing a much stronger correlation during EC (*r* = 0.77*, p <* 10*^−^*^9^) than EO (*r* = 0.39*, p* = 0.009). This is in line with our hypothesis that externally driven input during EO overrides the dynamics originating from local anatomy. Despite these differences, the distribution of dynamical dissimilarities is statistically indistinguishable between EC and EO (t-test in Figure 4c). Therefore, the dynamics during both EC and EO are different across areas (confirmed by MANOVA tests in Figure S8a), yet only the EC dynamics strongly correlate with the anatomy.

To fully understand the changes to dynamics caused by behaviour we also compared the univariate distributions of the spiking statistics between EC and EO (Figure S8b,c). We found the biggest changes in the average cross-correlation log(CC_avg_) especially for V1^L^, V4^L^ and M1*/*PMd^N&E^ (Figure S8c,d), in agreement with previous reports^36,40,41^.

### Autocorrelation timescales also reflect hierarchy

Autocorrelation timescales are a common metric for cortical hierarchy^5,18^. Previous studies have shown^5,18^ that sensory areas have shorter timescales, whereas higher areas have longer timescales. Our dynamical dissimilarity does not include the autocorrelation timescales, because these are notoriously difficult to estimate from limited data^37,38^ (see Figure S4). To overcome this problem, we estimated the timescales using longer data segments (t_segment_ = 300 s). We fitted an exponential function to the autocorrelation function (Figure 5a, see sample fits in Figure S9), analogous to previous studies^5,42^. See Autocorrelation timescales for a detailed description of the timescale estimation.

**Figure 5:**
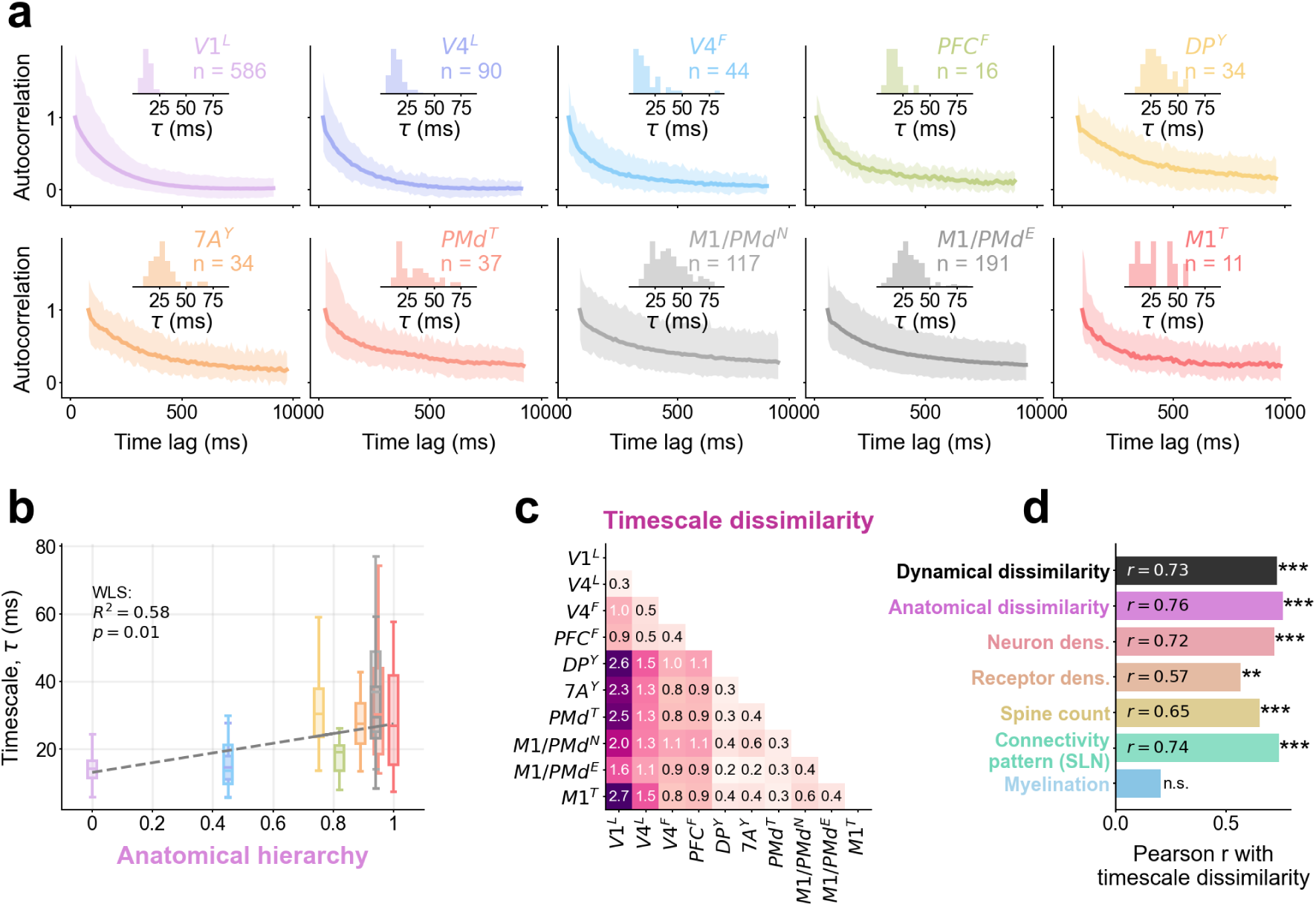
Autocorrelation timescales vary across the cerebral cortex and correlate with the anatomical hierarchy. **a)** Autocorrelation function for single neurons with an exponential fit for each area. The mean autocorrelation across neurons (dark line) and 5th to 95th percentile (shading) is shown. Insets show the distribution of the estimated timescales and the number of single neurons with a valid fit (*R*^2^ *>* 0.8). **b)** Boxplot of the estimated timescales for each area compared to their position in the anatomical hierarchy. WLS fit, its *R*^2^ and t-test p-value are shown. **c)** Timescale dissimilarity: Wasserstein distance between the timescale distributions. **d)** Pearson correlation coefficient between the timescale dissimilarity and other metrics: dynamical dissimilarity, anatomical dissimilarity, and inter-area differences in all the individual anatomical markers. Stars indicate result of t-test: *p >* 0.05*/k* (n.s.), 0.05*/k > p >* 0.01*/k* (*), 0.01*/k > p >* 0.001*/k* (**), *p <* 0.001*/k* (***). Bonferroni corrected for multiple testing (*k* = 61, also including the tests from Figure 3d).

As expected, we observed shorter timescales in sensory areas and longer timescales in motor areas (Figure 5a,b). Indeed, the timescales have a positive correlation with the anatomical hierarchy (Figure 5b). To compare the timescales to the other spiking statistics we measured the timescale dissimilarity, by taking the Wasserstein distance between the timescale distributions of every pair of areas (Figure 5c). We found significant correlations between the timescale dissimilarity and the other hierarchy markers (Figure 5d). As with the other spiking statistics (Figure 3d), timescale dissimilarities did not correlate significantly with myelination differences (Figure 5d). The correlation between the anatomical and timescale dissimilarities is very similar (Pearson’s *r* = 0.76, Figure 5d) to the correlation between the anatomical and dynamical dissimilarities (Pearson’s *r* = 0.74, Figure 3c). These results suggest that the timescales and the other spiking statistics are comparably robust indicators of the cortical hierarchy; but the spiking statistics can be measured faster and more robustly using far less data than the timescales (Figure S4).

## Discussion

This work has two main findings. First, we show that single-neuron spontaneous spiking statistics form a unique fingerprint of cortical areas, but only when considering multiple spiking statistics in combination. Second, our results reveal a strong correlation between the spontaneous spiking statistics and anatomical markers of cortical hierarchy, suggesting that the anatomical differences along the cortical hierarchy are imprinted within the spontaneous spiking activity, especially when the eyes are closed.

Our findings uncover rich and varied spiking statistics across different cortical areas, in agreement with previous reports^5,6,23^. Yet, the firing rate (FR) distributions were very similar across all cortical areas (Figure 2a). When considered alone, the FR distribution did not correlate with any of the anatomical markers of the cortical hierarchy (Figure 3d). This suggests that the FR distribution from spontaneous activity is not predominantly shaped by the local structure of the cerebral cortex, even though the FR is known to vary in an area-specific way during task-related activity.

Furthermore, we found no significant correlation between the myelination of cortical areas and their dynamics (Figure 3d and Figure 5d). This is surprising, since myelination is known to correlate with the other anatomical markers of the cortical hierarchy in macaques^4^ and myelination has also been shown to correlate with autocorrelation timescales in human^18,30^. This discrepancy may be explained in part by the fact that both Gao et al.^18^ and Cusinato et al.^30^ included data from more areas than our study. Furthermore, the common myelination proxy used here (T1w/T2w MRI signal) can include traversing axons that do not connect to, nor originate from, the measured area, thus introducing noise into this marker. This could cause a strong dependence of the correlation on the set of areas for which data are available, especially when the number of areas is small. In combination, our limited sample size and the observation uncertainty could explain the discrepancy. Additional spiking data and new direct measurements of myelination (from histology or polarized light imaging^43^) could help clarify this inconsistency.

Topographic connectivity between areas is ubiquitous in the brain^44^, such as retinotopic connectivity in visual areas^45^ or somatotopic connectivity in motor and somatosensory areas^46^. Cortical sites sharing more direct connections are known to have more similar spiking dynamics^45^. However, it was not possible to take these topographic connections into account for this study.

The hierarchical orders derived from dynamical and anatomical dissimilarities have a few notable differences (Figure 3e). The dynamical hierarchy positions of *M* 1*^T^* and *PMd^T^* are much lower than expected from the anatomical hierarchy. The reason for this could be that these two datasets include spiking activity from all cortical layers, whereas the other recordings contain data from a single layer. Since we did not have spiking data from all layers for most areas, we could not perform a layer-wise analysis.

The observed differences in spiking statistics between areas depend on both global influences from other brain areas and characteristics of each individual area. However, it is not possible to fully disentangle the global and local contributions *in vivo*. To prevent global stimulus-specific activity from dominating the dynamics, we study spontaneous activity, which should be less biased toward external stimuli than task-driven activity. We also studied eyes-open and eyes-closed activity separately (Figure 4), revealing a stronger correlation between the dynamical and anatomical dissimilarities with the eyes closed. As the monkeys were sitting in a dimly lit room, they received at least some visual input in the eyes-open condition. Another potential difference between eyes-open and eyes-closed conditions, even in the absence of visual stimuli, is the expectation of visual input when the eyes are open^36^. External stimuli or the readiness for visual input in the eyes-open condition may suppress the differences caused by the local structural differences between areas. If this is the case, our results suggest that cortical activity with substantial inter-area interactions is less hierarchical than that without. To eliminate the global component completely and study local dynamics, future work could employ the approach and metrics introduced in the current study on *in vitro* brain slice activity.

The hierarchical organisation of timescales has recently come into question with contradicting results in mice^7,19^. However, timescales in spiking activity have been reported to vary along the cortical hierarchy in the macaque brain^5,26^. As expected, our analysis found a significant correlation between the anatomical markers of the hierarchy and timescales in macaques, confirming previous findings. Interestingly, Cusinato et al.^30^ report large changes to the hierarchy of timescales between wakefulness and different sleep stages from human intracortical EEG recordings. Particularly, they report that timescales estimated from gamma-band signals (30-80 Hz) exhibit a hierarchy in sleep, but not during wakefulness, similar to our observations for macaque dynamical dissimilarities between EO and EC (Figure 4). We could not test whether the timescales change with behaviour in our data, since the periods of EO and EC are relatively short and do not allow for reliable timescale estimates (Figure S4). All our results focus on the pairwise differences between areas, e.g., the dynamical dissimilarity between V1 and M1 compared to the receptor density ratio between V1 and M1. However, the values for each area could carry similar information, e.g., the mean LvR in V1 compared to the mean receptor density in V1. Comparing the anatomical markers against the mean spiking statistics in each area (Figure S7) reveals some correlations, but most correlations are not statistically significant. Notably, no single statistic significantly correlates with all the anatomical markers. Thus, inter-area differences in spiking statistics more robustly reflect hierarchical patterns.

Comparing brain dynamics usually involves some metric in high dimensions, such as representational similarity via cross-correlation^47^, canonical correlation analysis^48^, hyperalignment^49^ or some shape metric^50^ (e.g., Procrustes distance). All these approaches are usually applied to spatio-temporal activity data. In contrast, our work explores an alternative way of comparing brain activity data, collapsing the time component and focusing on the space of a few relevant spiking statistics (firing rate, inter-spike interval variability, cross-correlation). One strength of our approach is its ability to compare heterogeneous datasets with varying electrode counts, recording duration, and experimental setup. Despite the heterogeneous data, the spiking statistics reliably distinguish cortical areas from each other (Figure 2) and reveal the link between their activity and the anatomical hierarchy (Figure 3).

Moreover, our approach is well-suited for data-to-model comparisons, which are usually challenging due to the vast subsampling in brain recordings. The distribution of multivariate spiking statistics is unique for each brain area (Figure 2), which could be used as a target distribution for parameter inference in spiking neuron models. The spiking statistics presented here could also be used to test the biological realism of large-scale neuronal network models^51,52^. One such study^52^ showed that introducing clustered connectivity in cortical models leads to similar FR distributions across areas.

While this study focused on areas within the visuomotor processing stream, our methods could readily be applied to other brain areas, such as those in the auditory and somatosensory processing streams. This could potentially uncover analogies and key differences between sensory modalities.

All in all, our findings reveal a fundamental link between cortical structure and spontaneous dynamics along the cortical hierarchy.

## Materials and methods

### Electrophysiological data collection of spontaneous activity

All data were collected from the cerebral cortex of macaque monkeys (*Macaca Mulatta*, N = 6) in the resting state, reflecting spontaneous cortical activity. The macaques were sitting in a dim-lit room, with no particular task, while the continuous activity from the cerebral cortex was recorded. The relevant behaviour in each experiment was also tracked using video recordings and/or eye tracking systems.

**Table 1:** Summary of subjects and recordings included in this study. Identity of the subject, cortical area, number of sessions, number of simultaneously recorded electrode sites in each session, and number of detected single-unit activities (SUA) are indicated.

| Subject | Area | # Sessions | # Contacts | # SUA |
| --- | --- | --- | --- | --- |
| L | V1 | 1 | 896 | 239 |
|  | V4 | 1 | 128 | 81 |
| F | V4 | 59 | 4 | 375 |
|  | PFC | 59 | 4 | 209 |
| Y | DP | 1 | 36 | 39 |
|  | 7A | 1 | 36 | 47 |
| T | M1 | 7 | 24 | 89 |
|  | PMd | 11 | 24 | 57 |
| N | M1/PMd | 2 | 96 | 310 |
| E | M1/PMd | 2 | 96 | 277 |

### Data from macaque L

The data from macaque L were recorded from visual areas V1 and V4 (N=1 subject, N=1 session of 20 min duration). Chronic recordings were made using 16 8×8 electrode Utah arrays (Blackrock microsystems), two of them in visual area V4 and the rest in the primary visual cortex (V1), with a total of 1024 electrodes. The electrodes were 1.5 mm long; thus, the recordings were likely made from the deeper layers L5 and L6. A full description of the experimental setup and the data collection and preprocessing has already been published^9^.

Pupil position and diameter data were collected using an infrared camera to determine the direction of gaze and eye closure of the macaques. In addition to the spontaneous activity recordings, a visual response task was also performed. The visual response data were used to calculate the signal-to-noise ratio (SNR) of each electrode and all electrodes with an SNR lower than 2 were excluded from further analysis.

The raw data were spike-sorted using a semi-automatic workflow, as described in a previous publication^36^.

All experimental and surgical procedures for macaque L complied with the NIH Guide for Care and Use of Laboratory Animals, and were approved by the institutional animal care and use committee of the Royal Netherlands Academy of Arts and Sciences (approval number AVD-8010020171046).

### Data from macaque F

The data from macaque F were recorded from visual area V4 and prefrontal cortex (PFC) including Brodmann areas 8, 45A, and 46 below the principal sulcus (N=1 subject, N=59 sessions of 5 min duration). Acute recordings were made using the Omniplex system (Plexon Inc., Dallas, TX). On a given day, up to four glass-coated tungsten microelectrodes (impedance 1-1.5 MOhm, Alpha-Omega Engineering) were positioned through a grid system over each area (PFC and V4) and were advanced through the dura by a four-channel microdrive system (NAN Instruments, Nazareth, Israel). Signals were filtered between 300 Hz (4-pole high pass Bessel) and 8 kHz (2-pole high pass Bessel), amplified, and digitized at 40 kHz to obtain spike data. Recordings were carried out within the first 1-1.5mm after encountering the first activity; thus, the majority of data were obtained from the middle and superficial layers (L2/3/4). The spontaneous activity was recorded while the macaque was not performing any particular task. A full description of the experimental setup and the data collection and preprocessing—for the accompanying experiment in the same subject—has already been published^12^.

Eye position was monitored by an infrared-based tracking system (ETL-200, ISCAN Inc., Woburn, MA) and was sampled at 240 Hz. The x and y position were recorded and used for behavioural segmentation into eyes-open and eyes-closed epochs. Offline spike sorting was carried out manually using template matching in Offline Sorter (Plexon Inc). Four to ten clean single units were identified per area and session.

All experimental and surgical procedures for macaque F were approved by the Institutional Experimental Protocol Evaluation Committee (Approval Number 108596/30-5-2017).

### Data from macaque Y

The data from macaque Y were obtained from two chronic Utah arrays (Blackrock microsystems), one in dorsal prelunate cortex (area DP) and another one in parietal area 7A, for a total of 72 electrodes (N=1 subject, N=2 sessions of 10 min duration). The recording apparatus is described elsewhere^11^. The electrodes were 1 mm long, thus recording from the central layers, likely layer 4. The recording system recorded the electric potential at each electrode with a sampling rate of 30 kHz. Spontaneous activity data from the same experiment have previously been analysed and published^36^.

The raw data were spike-sorted for this study using a semi-automatic workflow using spikeinterface^53^. After the automatic sorting the quality of the waveform clusters was manually evaluated and only clean single-unit activity (SUA) waveforms were included in this study.

All experimental and surgical procedures for macaque Y were approved by the local (INT, Marseille) ethical committee (C2EA 71; authorization Apafis#13894-2018030217116218v4) and conformed to the European and French government regulations.

### Data from macaque T

The data from macaque T were recorded from premotor cortex (PMd) and primary motor cortex (M1) (N=1 subject, N=11 sessions of 15 min duration). Acute recordings were made with laminar probes (Plexon and Alpha Omega, 24 contacts, 100 and 200 µm pitch). The laminar probes enabled recording from across the cortical layers. The 200 µm pitch probes could record all layers simultaneously, while the 100 µm pitch probes did not span the entirety of the cortical gray matter. The motor cortex gray matter is known to be approximately 3.5 mm thick^54^, with the superficial and deep layers roughly equal in thickness. A full description of the experimental setup and the data collection and preprocessing—for the accompanying experiment in the same subject—has already been published^55^.

The raw data was spike sorted offline. Spike sorting identified 5-13 clean single units per probe and session. In addition to the spiking data, surface Electromyography (EMG) of the contralateral deltoid muscle, the heart rate with an ear clip, and a video of the macaque behaviour were recorded. In all the behavioural videos the screen LEDs were used to send a 1 s long blink every minute that can be used to realign the video with the neural recordings. We performed a video-based segmentation into behavioural epochs: eyes-open, eyes-closed, and movement periods.

Care and treatment of macaque T during all stages of the experiments conformed to the European Commission Regulations (Directive 2010/63/EU on the protection of animals used for scientific purposes) applied to French laws (decision of the 1st of February 2013). The experimental protocol was evaluated by the local Ethics Committee (CEEA 071) and carried out in a licensed institution (B1301404) under the authorization 03383.02 delivered by the French Ministry of High Education and Research.

### Data from macaques N & E

The data from macaques N & E were recorded from the interface between premotor (PMd) and primary motor (M1) cortex (N=2 subjects, N=2 sessions of 15-20 min duration per subject). The recordings were chronic using one 10×10 electrode Utah array (Blackrock microsystems).

A full description of the experimental setup and the data collection and preprocessing—for the accompanying experiment in the same subject—has already been published^10^. An extensive analysis of the spontaneous data has also been published^40^.

In addition to the registration of brain activity, the monkey’s behaviour was video recorded and synchronized with the electrophysiology recording.

All experimental and surgical procedures for macaque N & E were approved by the local (INT, Marseille) ethical committee (C2EA 71; authorization A1/10/12) and conformed to the European and French government regulations.

### Spiking data preprocessing

To ensure reliable analysis, several preprocessing steps common to all datasets were taken.

The data were sliced into segments of 5-second length. The segments were fully contained within one behavioural epoch: eyes-open, eyes-closed, or movement. The behaviour was estimated from video-tracking, eye-tracking and/or muscle activity, depending on the available setup in each experiment. Spontaneous spiking activity is known to vary strongly between different behaviours^36,40^. Including samples uniquely in one epoch ensured that major activity transients from behavioural transitions were excluded.

Furthermore, electrophysiology recordings are prone to artifacts from external noise and cross-talk between recording channels^31^. These artifacts can be detected as higher-than-chance synchronous activity at sampling resolution (30 kHz)^9,31^. Using the methods proposed by Oberste-Frielinghaus et al^31^, we estimated the excess synchrony and removed all units suspected to originate from common noise or cross-talk, which would especially affect the correlation statistics in our analysis.

### Statistics of spiking neuronal data

The experimental spike trains were sliced into 5-second segments. We computed a number of statistics for each one of these data samples and constructed a multivariate cloud of neuronal spiking statistics. These multivariate spiking statistics are used to characterise a given area and assess the similarities across areas.

We used the following spike train statistics:

- Mean firing rate (FR), given as the total number of spikes divided by the duration of the recording sample. Since the FR has a right-skewed distribution, we take its logarithm throughout this paper. The distribution of log(FR) better resembles a normal distribution.
- Revised local variation of the inter-spike intervals (LvR)^6^,

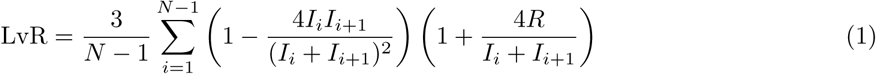

where *N* is the total number of spikes, *I_i_* is the *i*th inter-spike interval, and *R* = 5 ms is the refractoriness constant. A value of LvR = 0 corresponds to perfectly rhythmic spiking, whereas LvR = 1 corresponds to the irregularity of a Poisson point process.

- The average of the Pearson cross-correlation of the spike counts (bin width = 10 ms) with all other simultaneously recorded neurons (CC_avg_). This metric quantifies whether the measured neuron coordinates its activity with many of the other neurons (i.e., if it is a chorister) or if it is spiking independently (i.e., is a soloist)^32^.
- The standard deviation of the Pearson cross-correlation of the spike counts (bin width = 10 ms) with all other simultaneously recorded neurons (CC_std_).

Further statistics were considered but not used in our analysis, because they were strongly correlated with other statistics. If strongly correlated statistics were introduced, some spiking properties would be over-represented. For example, the coefficient of variation (Cv, Cv2)^56^ and local variation (Lv)^23^ of the inter-spike intervals are excluded due to their similarity to the LvR. Introducing all of Cv, Cv2, Lv and LvR would significantly increase the representation of the inter-spike interval variation, without a large increase in the explained variance. Another statistic that was considered was the spike-triggered population response (stPR)^32^, but it was not included due to its similarity with CC_avg_.

To exclude outliers, potential artifacts, or numerical issues, the statistics were bounded to the following empirically determined ranges: FR *>* 1 spikes*/*s, LvR ∈ [0, 3], CC_avg_ *>* 10*^−^*^4^, and CC_std_ *>* 0.01. We ensured that the selection of these ranges did not distort the underlying distributions or truncate any long tail, instead excluding only outliers. See Figure S1–S3 for the full distribution of included spiking statistics for all datasets.

All metrics were computed using their implementation in the elephant toolbox^57^, within the NetworkUnit reproducible testing framework^58^.

### Statistical testing with ANOVA and MANOVA

For all ANOVA and MANOVA tests, we used a fixed number of points from the multivariate spiking statistics (*N* = 500, randomly sampled), since the tests are robust to the normality assumption if the samples are large (generally *N >* 30) and robust to the assumption of equal (co-)variance (homoscedasticity) if the sample sizes are equal^59^. The significance level was Bonferroni corrected for multiple testing (*k* = 90 experiment pairs · 7 test types = 630), such that *α* = 0.05*/k* = 7.9 · 10*^−^*^5^.

### Dynamical dissimilarity for comparison of multivariate and univariate distributions

To compare the distributions of spiking statistics to each other we used the Wasserstein distance (WS). The WS is defined as an optimal transport distance between two observed distributions. Each multivariate spiking statistics cloud is an *N* × *M* matrix of *N* samples (spike train segments) and *M* spiking statistics. The compared distributions must have the same number of spiking statistics *M*, but can have different sample sizes *N* . To avoid potential sampling bias, we used equal sample sizes *N* in our analysis.

We normalized (z-scored) the statistics, to ensure that all metrics *M* were equally weighted for the distance measurement. We assigned equal weights *w* to each neuron within each distribution, such that *^N^ w* = 1. Finally, we searched for the optimal transport between the two distributions. Note that it is not necessarily a one-to-one map since the weight from one node can be redistributed to various nodes. The Wasserstein distance is the work normalized by the transported weight,

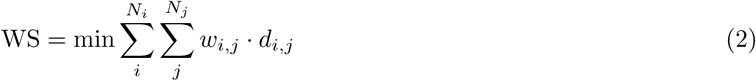

where *N_i_*, *N_j_* are the number of samples in the two compared spiking statistics clouds, *w_i,j_* is the weight transported between points *i* and *j*, and *d_i,j_* is the Euclidean distance between them. The weight normalization 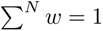 ensures that also 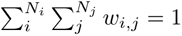. To find the optimal transport configuration we used a dedicated simplex algorithm implemented in OpenCV^60^.

We studied both the multivariate and univariate distributions using the WS distance.

### Anatomical data for hierarchical characterization

The anatomical data for hierarchical characterization of the cortical areas was collated from various published sources in the literature.

Neuron densities were estimated from Nissl- and NeuN-stained histological slices of macaque^13,61^. Since the NeuN method tends to underestimate neuron densities^62^, we linearly scaled NeuN estimates to match the Nissl-stain estimates of neuron density^14,51^, based on areas for which both estimates were available. This provided appropriately scaled data for those areas for which only NeuN staining was available.

Receptor densities were measured via *in vitro* receptor autoradiography for 14 different receptor types using receptor-specific protocols^4^. The 14 receptor types included the following; Glutamate: AMPA, kainate and NMDA; GABA: GABA_A_, GABA_A_*/*BZ and GABA_B_; acetylcholine: M_1_, M_2_ and M_3_; serotonin: 5 − HT_1A_ and 5 − HT_2A_; noradrenaline: *α*_1_ and *α*_2_; and dopamine: D_1_. We used the average receptor density as a proxy for the cortical hierarchy (Figure 3), and also report the correlations for each individual receptor (Figure S5).

Spine counts for several cortical areas in the macaque were previously described by Elston^16^, later mapped to a common parcellation and openly shared by Froudist-Walsh et al.^15^.

The ratio of T1-weighted to T2-weighted (T1w/T2w) MRI signal is a proposed marker of myelination in the cortical gray matter. T1w/T2w was measured by Donahue et al.^17^, later mapped to a common parcellation and openly shared by Froudist-Walsh et al.^4^.

The hierarchical level of cortical areas is classically defined from the structural connectivity between cortical areas^1^. In particular, the laminar connectivity patterns have been shown to follow a hierarchical organisation^2^. Here, we use the hierarchical levels estimated and shared by Froudist-Walsh et al.^15^.

Many anatomical markers are known to have heavy-tailed distributions, such as the lognormal distribution^14,63^. Indeed, the neuron density, mean receptor density, spine count, and myelination have right-skewed distributions; thus, we took their base-10 logarithm. The logarithm of a lognormal distribution is a normal distribution, which is better suited for statistical testing and correlation measurements.

For this work, we calculated the anatomical dissimilarity by taking the Euclidean distance between the 5D points reflecting the neuron density, mean receptor density, spine count, and hierarchical levels from connectivity patterns. The individual anatomical markers were first normalized between 0 and 1.

### Estimation of hierarchies from dynamical and anatomical dissimilarities

To estimate the hierarchical position of each area from the dynamical or anatomical dissimilarities, we embed them into a 1-dimensional line using Isomap (*k* = 3 neighbours). The resulting embedding is normalized between 0 and 1. For the embedding, we use the full symmetric dissimilarity matrices, not just the lower triangular matrices shown in Figure 3a,b.

Note that for the embedding of the anatomical dissimilarities, we excluded the myelination.

### Autocorrelation timescales

To estimate the autocorrelation timescale, we first estimated the firing rate of all neurons using a 10-ms bin size. Then, we z-scored the firing rate, calculated the autocorrelation by convolving the signal with itself, and normalized the autocorrelation by its maximum value. Finally, we fitted an exponential function to the autocorrelation using least-squares (scipy.optimize.curve fit). The exponential function is of the form:

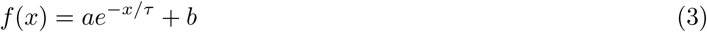

where *a* and *b* are arbitrary scaling constants and *τ* is the autocorrelation timescale. Following Fontanier et al.^42^, if there was an autocorrelation peak within the first 100-ms of time lag, we fit only starting from that peak and not from zero lag. We consider the neuron to lack a valid (measurable) *τ* when the exponential function fit does not converge or it has a coefficient of determination *R*^2^ ≤ 0.8.

For the main results (Figure 5) we split the spike trains of each neuron into 300 s data segments. One fit was calculated for each segment, without averaging across segments for the same neuron. Shorter data segments yielded only few valid measurements of *τ* (Figure S4). Example autocorrelation functions and fits with valid *τ* for a few randomly selected neurons are shown in Figure S9.

## Acknowledgements

We thank Jon Martinez Corral for proofreading the manuscript; Sarah Palmis for support with data collection from macaque T; Anno Kurth for helpful discussions about statistics and high-dimensional spaces; Nilanjana Nandi for helpful discussions about autocorrelation timescales; Ján Antolík and Karolína Korvasová for feedback on the manuscript about the interpretability of the results.

This work received funding from the Programme Johannes Amos Comenius (OP JAK) under the project ’MSCA Fellowships CZ - UK3’ (reg. n. CZ.02.01.01/00/22 010/0008220); the DFG Priority Program (SPP 2041 ”Computational Connectomics”) [S.J.van Albada: AL 2041/1-1]; the EU’s Horizon 2020 Framework Grant Agreement No. 785907 (HBP-SGA2) and No. 945539 (HBP-SGA3); the DFG (RTG 2416 ”MultiSenses-MultiScales”); the CNRS Multidisciplinary Exploratory Projects initiative; and a grant co-financed by Greece and the European Union (European Regional Development Fund) through the Operational Programme “Competitiveness Entrepreneurship Innovation 2014–2020” (to G.G. Gregoriou project MIS 5070462) in the context of the FLAG-ERA project PrimCorNet.

## Author contributions

Conceptualization AMG. Data collection XC (macaque L); SP, PS, GG (macaque F); BEK (macaque T). Spike sorting AMG (macaque L); AKJ (macaque Y); SP, PS, GG (macaque F); BEK (macaque T). Data curation and preprocessing AMG. Methodology AMG, RG, BEK, SvA. Software AMG, RG. Formal Analysis AMG. Visualization AMG. Writing - original draft AMG. Writing - review & editing all authors. Supervision SvA, BEK. Resources GG, TB, BEK, SvA. Funding acquisition AMG, GG, TB, BEK, SvA.

## Data and code availability

All data (Spike sorted activity recordings, anatomical data, measured statistics, dissimilarity metrics) and the code required to reproduce this analysis are publicly available (https://doi.org/10.5281/zenodo.17601400) under a Creative Commons 4.0 Attribution license (CC-BY 4.0).

## Supplementary figures

**Figure S1:**
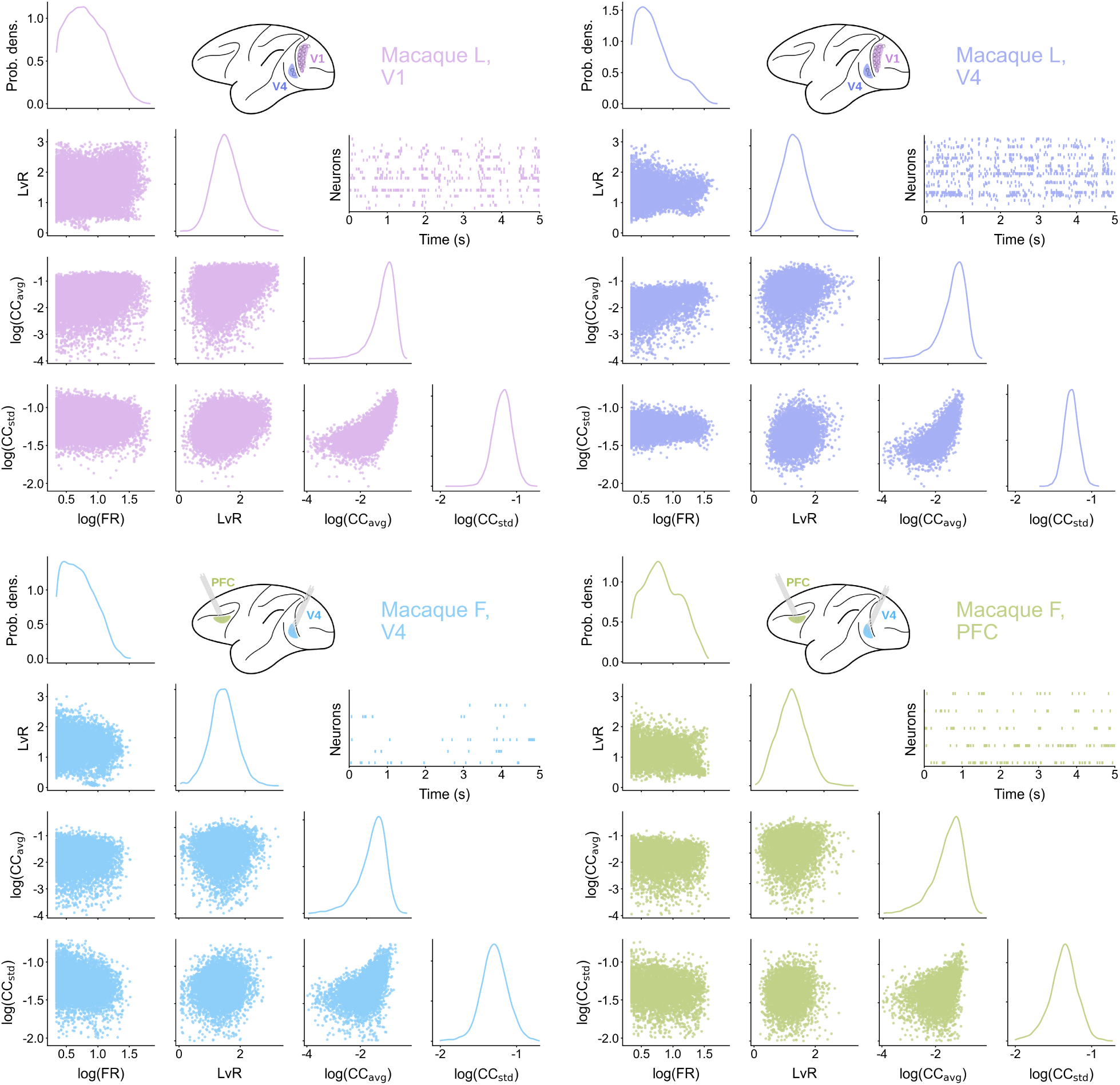
Multivariate spiking statistics for macaque L, areas V1 (top left) and V4 (top right), and macaque F, areas V4 (bottom left) and PFC (bottom right). Sample spike trains shown.

**Figure S2:**
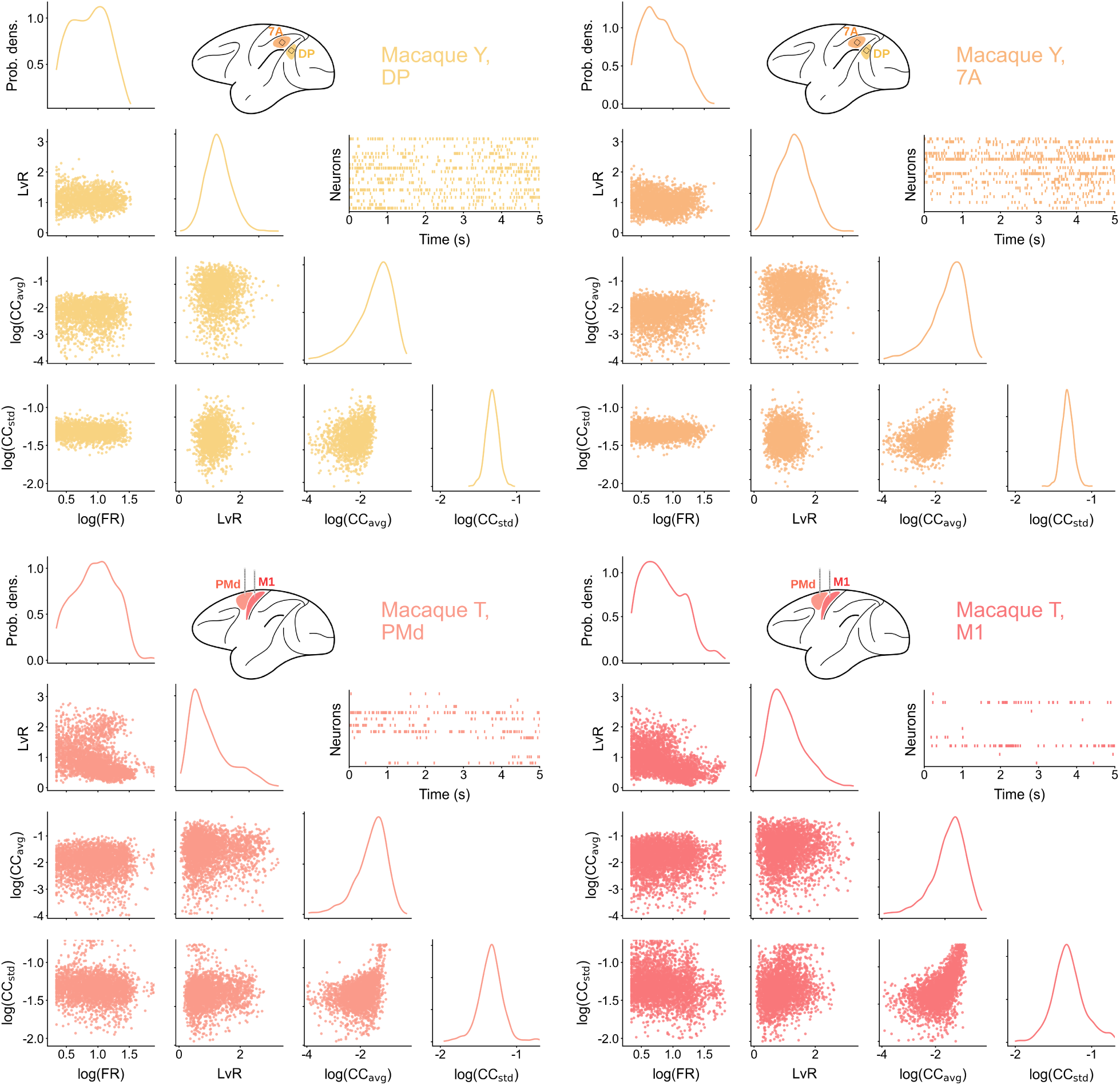
Multivariate spiking statistics for macaque Y, areas DP (top left) and 7A (top right), and macaque T, areas PMd (bottom left) and M1 (bottom right). Sample spike trains shown.

**Figure S3:**
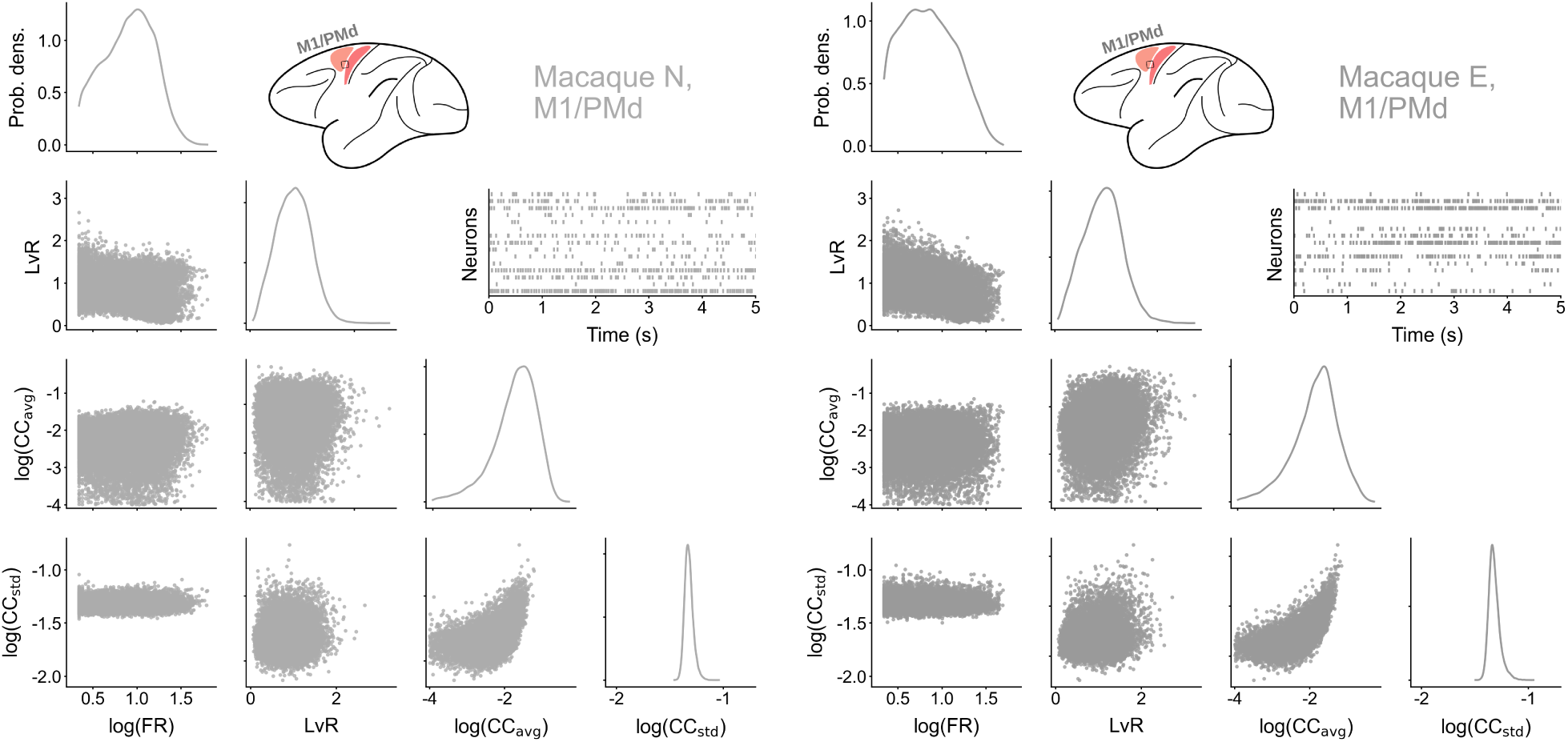
Multivariate spiking statistics for macaque N (left) and macaque E (right), area M1/PMd in both cases. Sample spike trains shown.

**Figure S4:**
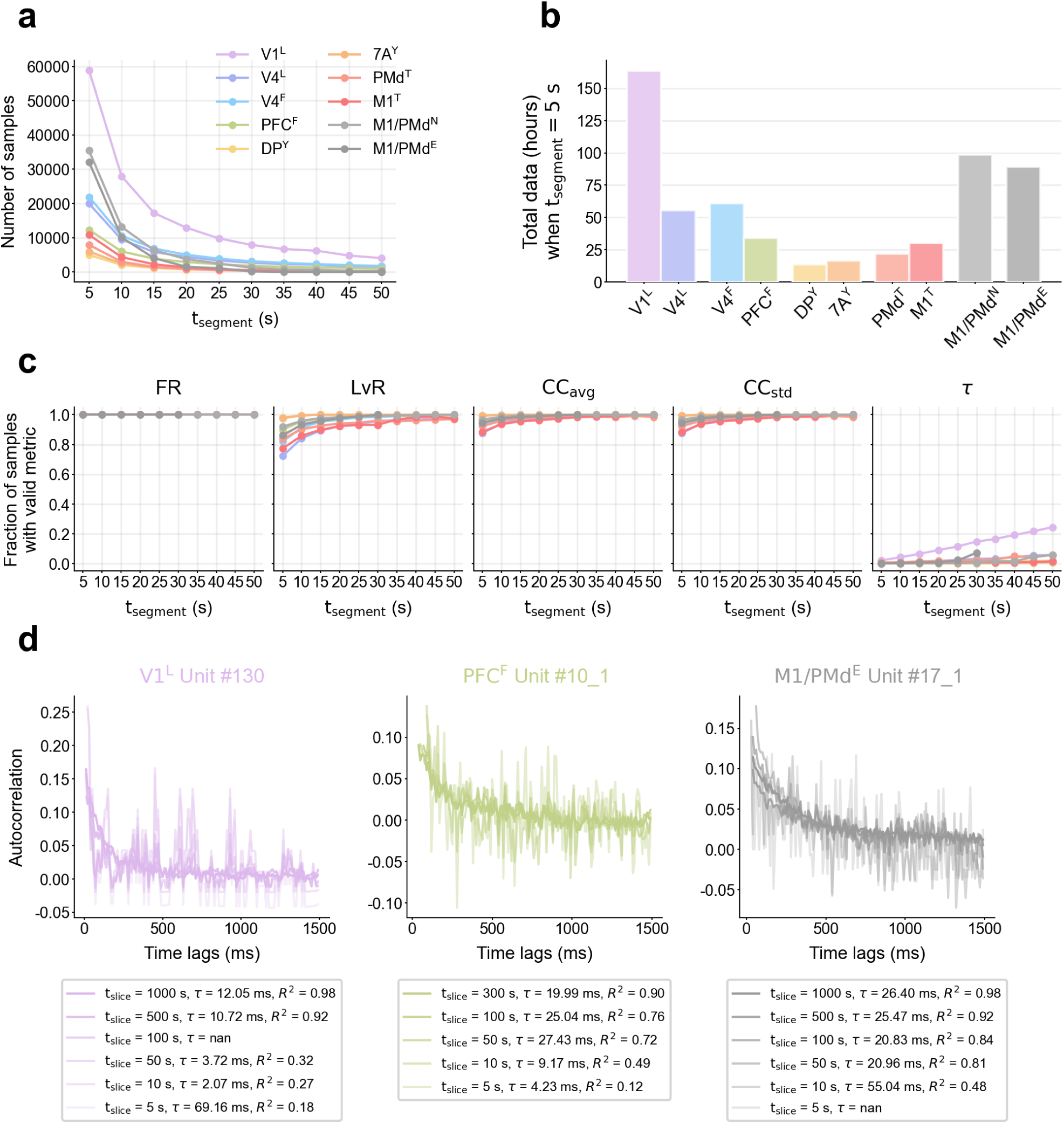
Estimation of autocorrelation timescales heavily depends on data length. **a)** Total number of samples when data is sliced at different lengths. Each segment is only allowed to contain one behavioral epoch. **b)** Total amount of available data (when t_segment_ = 5 s), given in hours. Note that segments of simultaneously recorded neurons are counted separately. **c)** Fraction of samples with a valid metric as a function of segment lengths t_segment_. FR is always well defined, even when no spikes are present. LvR cannot be calculated in segments with fewer than 3 spikes, which happens in less than 20% of samples. CC_avg_ and CC_std_ cannot be calculated if one or both segments have no spikes, which happens in less than 10% of samples. The autocorrelation timescale *τ* is only valid in a small fraction of the data samples (we consider *τ* invalid when the exponential function fit does not converge or it has *R*^2^ ≤ 0.8). Increasing t_segment_ drastically reduces the number of samples and for many, it is still not possible to estimate *τ* . **d)** Selected single-neuron examples where timescales fail to converge in short segments, but produce robust values for very long data segments.

**Figure S5:**
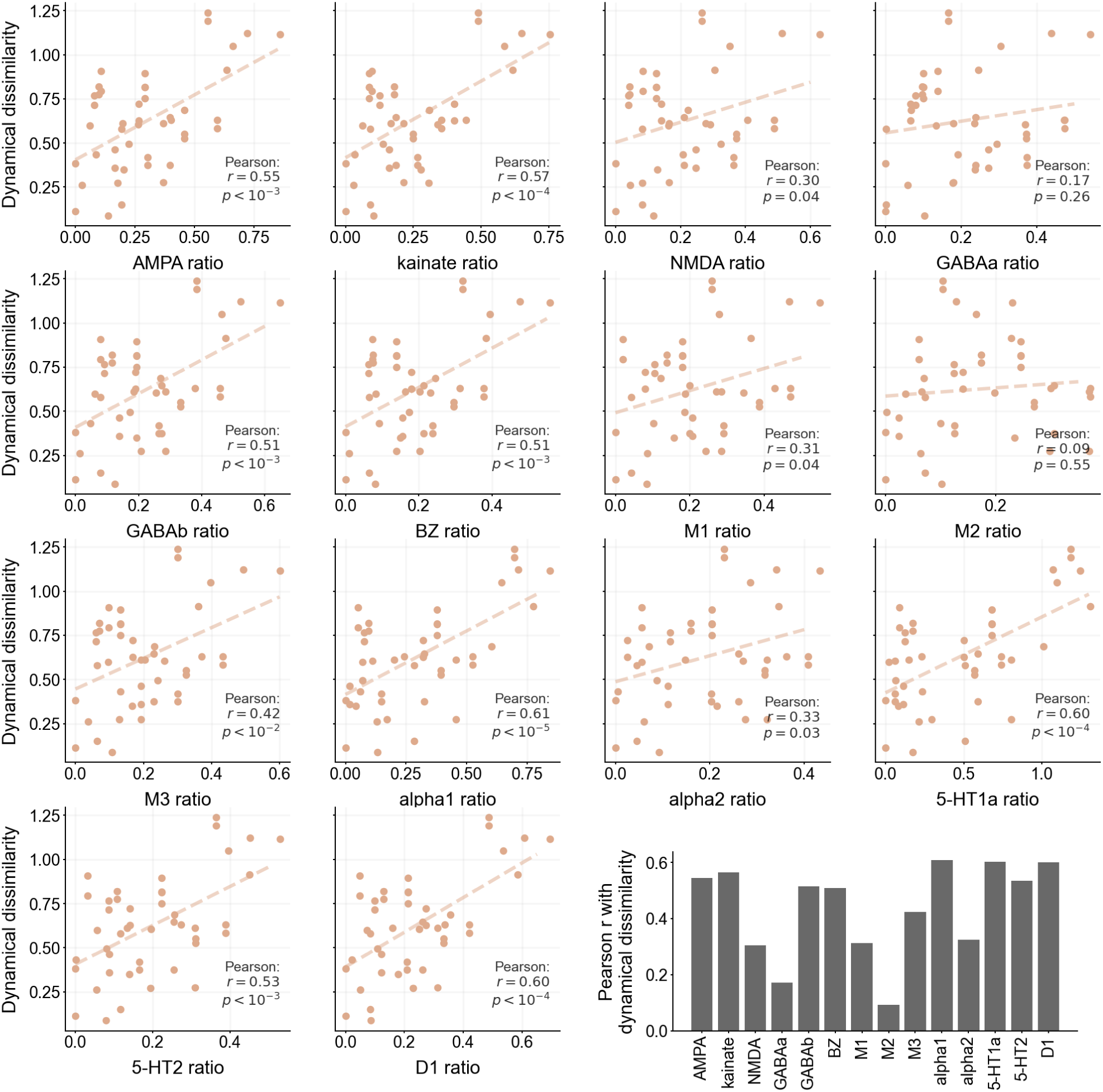
Correlation between the multivariate dynamical dissimilarity and the logarithm of the density ratio for all 14 receptors individually. Lower barplot summarizes the Pearson r coefficients. All receptors strongly correlate with the multivariate dynamical dissimilarity except GABAa and M2.

**Figure S6:**
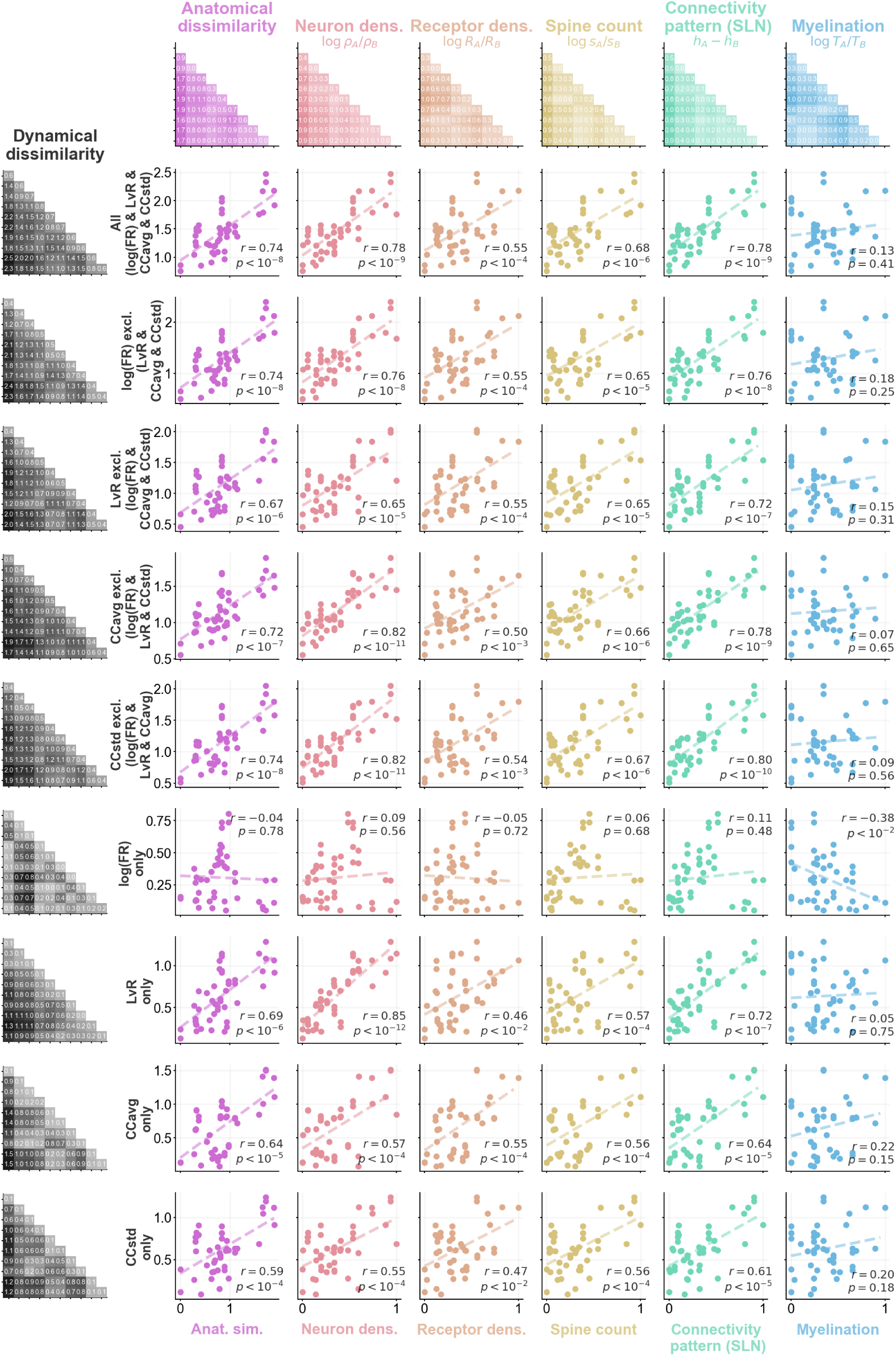
Detailed scatter plots corresponding to the correlations reported in Figure 3d. All the variants of dynamical dissimilarity (excluding one, or including only one metric) and all the individual anatomical differences shown. For each pair of variables, Pearson’s r and the associated p-value are shown, which correspond to the values from Figure 3d.

**Figure S7:**
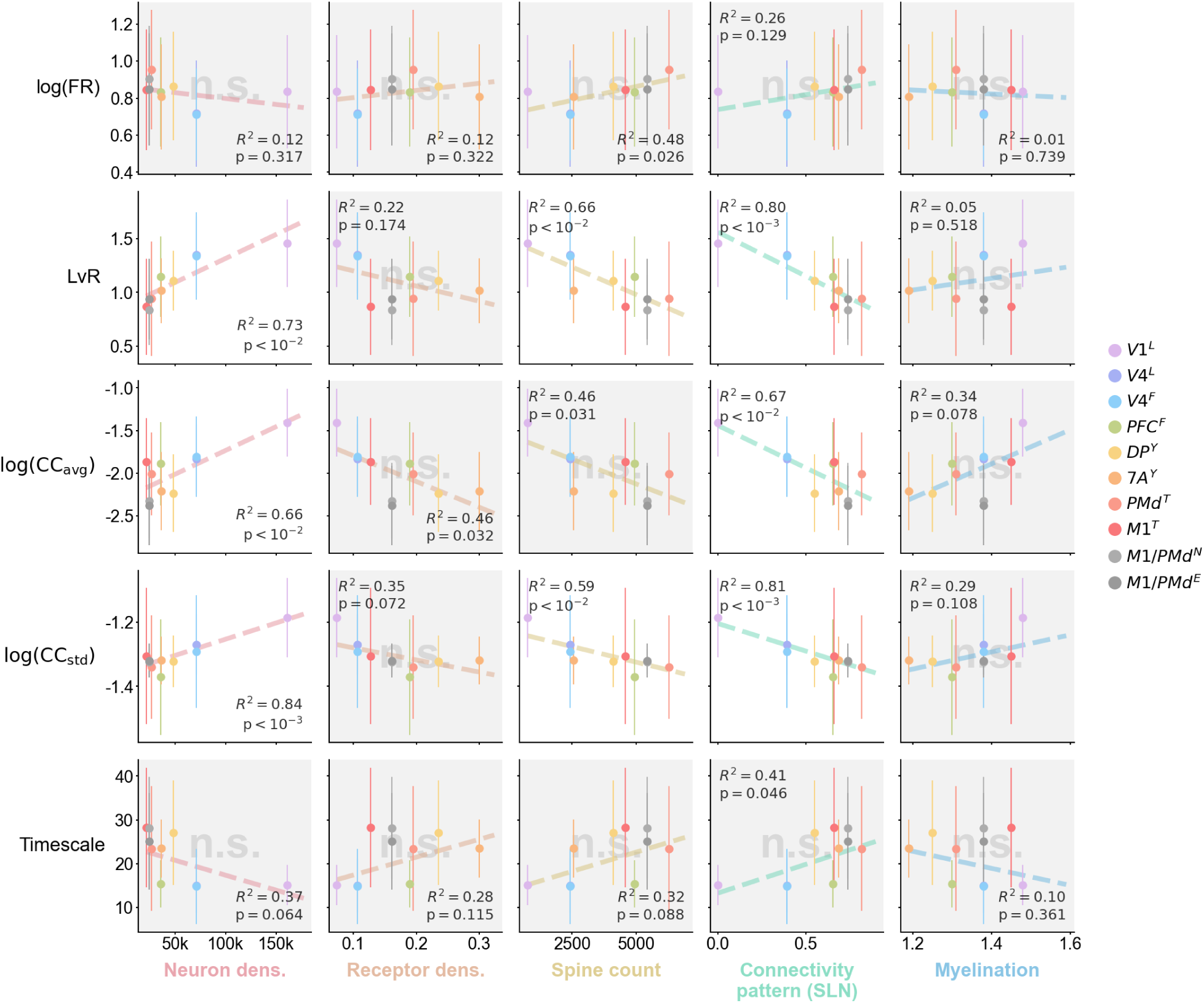
Correlation between the individual anatomical markers (x-axis) and the spiking statistics (y-axis). The error bars indicate the variance of the spiking statistics. A linear fit is performed via weighted least squares, where the weights are the inverse of the variance. R^2^ of the fit and the results of the corresponding t-test are shown. The correlation is considered non-significant (n.s.) if *p >* 0.05*/k*, with the Bonferroni correction of *k* = 5 tests per dataset. Non significant panels have gray shading and an ’n.s.’ watermark. No single spiking statistic correlates with all anatomical markers. Note that here we consider each area individually, whereas in Figure 3a–d and Figure 5c–d the areas are compared in a pairwise manner.

**Figure S8:**
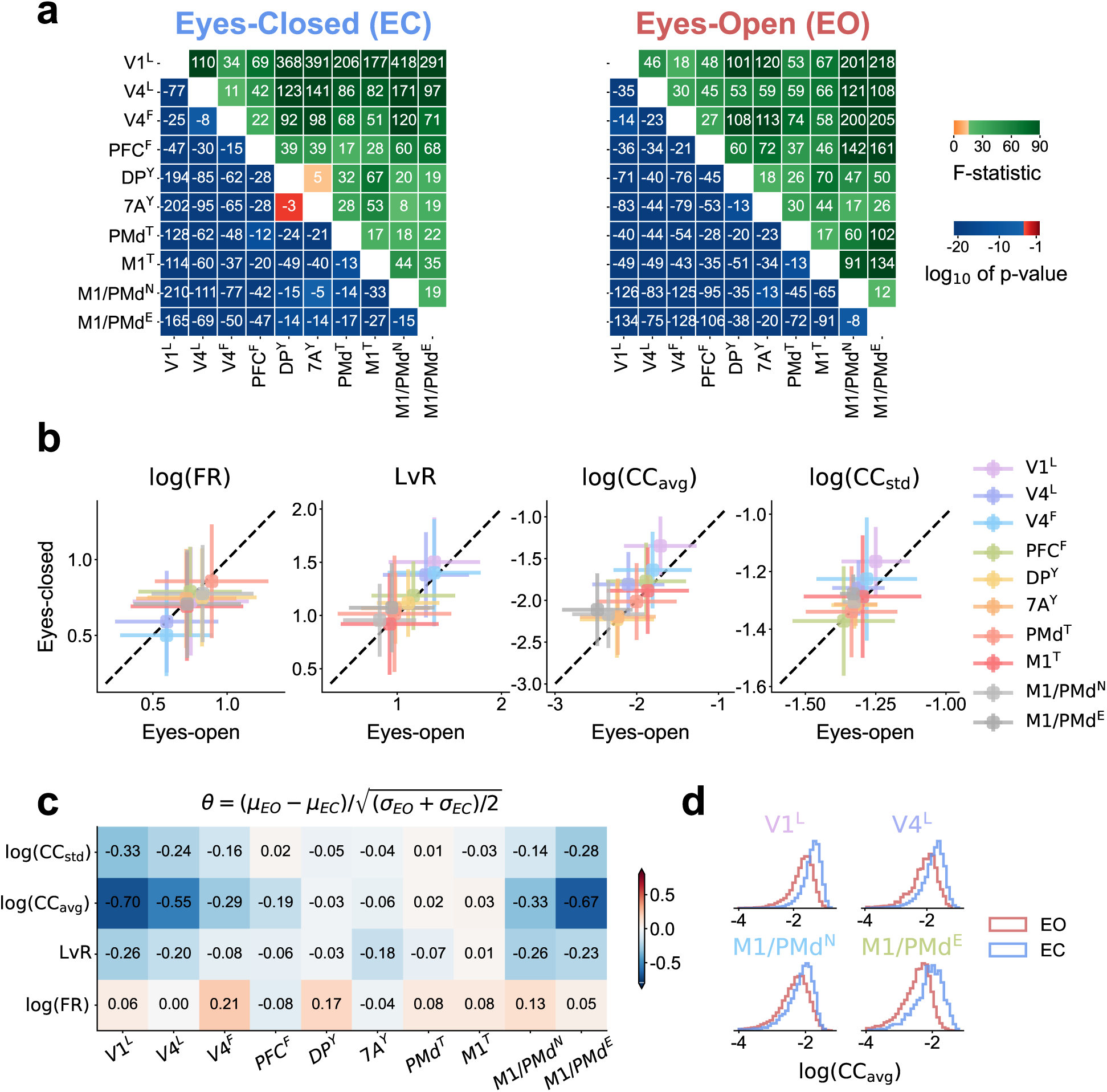
Supplementary analysis to the differences between eyes-open (EO) and eyes-closed (EC) conditions. **a** MANOVA test between all areas when including exclusively EC or EO data. Lower triangular entries show the base-10 logarithm of the p-values and the upper triangular entries show the F-statistic. Significance levels (*α* = 0.05*/k*) are corrected for multiple testing following the Bonferroni correction (*k* = 630 including all the tests from Figure 2). The significance level is color-coded, with red p-values (orange F-statistics) denoting non-significant results. The test concludes that nearly all area dynamics are different from each other both in EC and EO. **b** Comparison of mean and std (errorbar) between EO and EC for all spiking statistics. For cases where the distributions do not change between EO and EC the mean should lie along the diagonal; conversely significant changes diverge from the di<u>agonal.</u> **<u>c</u>** <u>Meas</u>urement of EO/EC differences for each spiking statistic and area. We report *θ* = (*µ_EO_* − *µ_EC_*)*/* (*σ_EO_* + *σ_EC_*)*/*2, which measures the changes to the mean in terms of the standard deviation. Most cases have small or even non-noticeable changes. **d** Sample distributions of log(CC_avg_) for the four areas with the largest differences between EO and EC.

**Figure S9:**
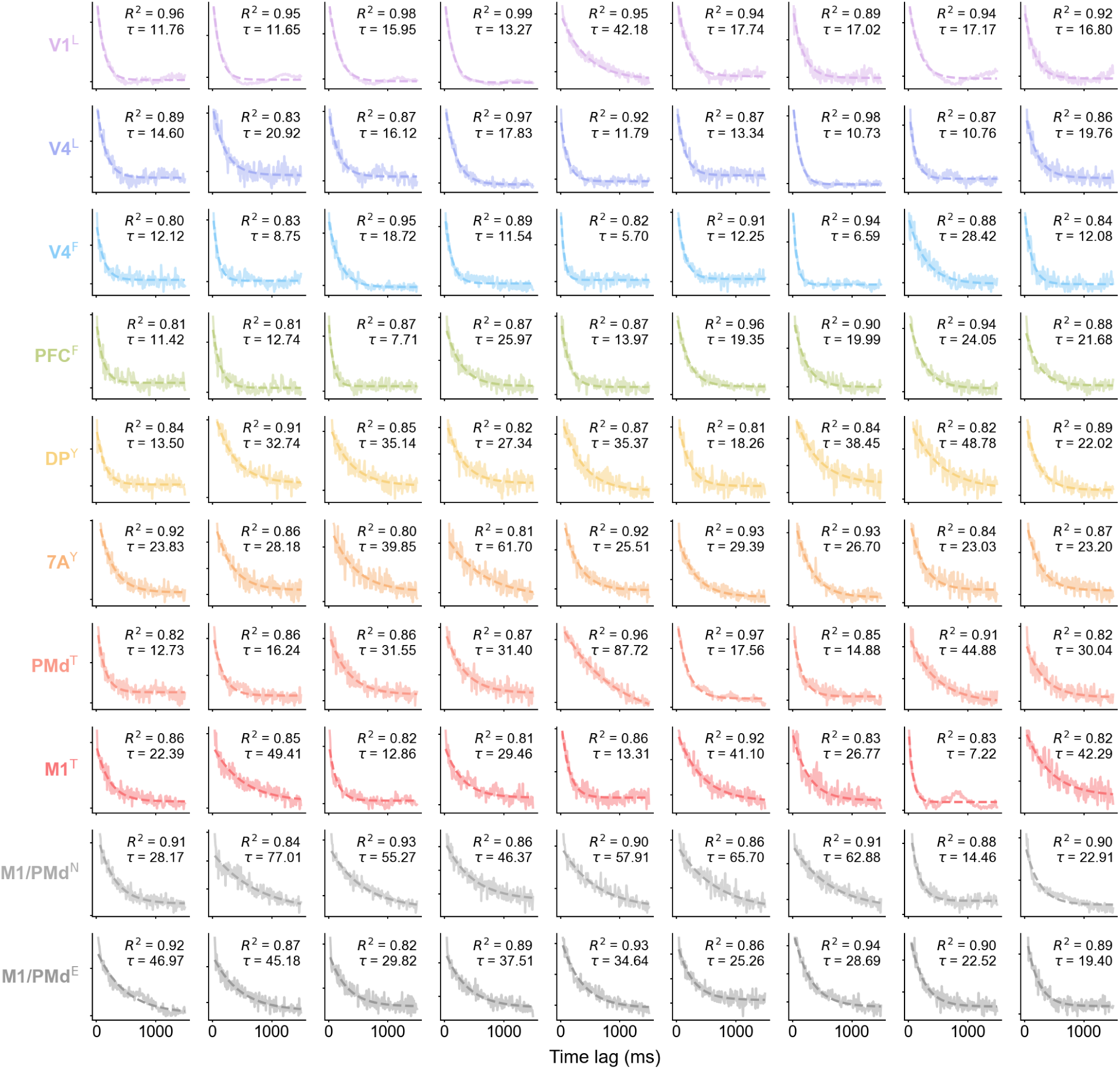
Example autocorrelation functions of single-neuron spike trains and the corresponding exponential fits (*R*^2^ and *τ* shown). The examples were randomly selected out of the neurons that had a good fit (*R*^2^ *>* 0.8).

